# TD2: finding protein coding regions in transcripts

**DOI:** 10.1101/2025.04.13.648579

**Authors:** Alan Mao, Hyun Joo Ji, Brian J. Haas, Steven L. Salzberg, Markus J. Sommer

**Author notes:** **Corresponding author** Markus J. Sommer.

## Abstract

The transcriptome encompasses all RNA transcripts in eukaryotic cells, orchestrating gene expression and regulating cellular function, development, and adaptation. Identifying open reading frames (ORFs) in transcripts is a critical step in transcriptome analysis. We introduce TD2, a new tool for *ab initio* annotation of protein-coding ORFs in transcripts. We find TD2 to be sensitive and precise when compared to other state-of-the-art tools in reference transcripts and transcriptome assemblies from a diverse array of eukaryotes.

TD2 is available at https://github.com/Markusjsommer/TD2. The project is open-source, developed in Python with PyTorch, and is freely available to all academic, government, and commercial users under the MIT license.

## Installation

TD2 can be installed with “*pip install TD2*” and is distributed via the Python Package Index (PyPI). TD2 may also be downloaded from https://github.com/Markusjsommer/TD2 and installed via “*pip install*.” from within the source directory.

### Step 1: finding candidate ORFs with TD2.LongOrfs

To identify candidate coding regions from a set of transcripts, TD2 identifies all translation initiation and termination codons, then generates a complete set of ORFs for each of the 6 reading frames per transcript. TD2.LongOrfs provides support for all 26 current NCBI translation tables, as well as alternative start codons for each table. The table number and use of alternative start codons can be specified in the command. TD2 is parallelized and multithreaded by default.

### Step 2: predicting protein coding regions with TD2.Predict

TD2 uses PSAURON ^1^, a pretrained machine learning model, to score protein coding regions. Candidate ORFs that receive a positive assessment from PSAURON, or are above a minimum length threshold, are retained in the final output. To increase sensitivity to known proteins, users may also supply TD2 with homology search results. Significant hits from blastp ^2^, HMMER3 ^3^, or MMseqs2 ^4^ cause TD2 to retain ORFs regardless of PSAURON score.

A few of the more impactful TD2 options are detailed here:

- Partial ORFs, those without both a start and stop codon, are retained by default. Users may restrict their analysis to complete ORFs with the *--complete-orfs-only* option. This option ensures all ORFs have both a start and stop codon and do not run off the edge of a transcript.
- TD2 will choose the single best ORF per transcript by default, similar to the *--single-best-only* option in TransDecoder. Users may turn this behavior off by using the *--all-good* flag, which will report all ORFs that pass PSAURON or length-based false discovery filters.
- The minimum in-frame PSAURON score required to retain an ORF is 0.50 by default. Users may change this cutoff with the *-P* option. The cutoff may range from 0 to 1, and a higher cutoff results in less sensitive and more precise predictions.
- TD2 may be run in precise mode, which adjusts parameters to increase precision while reducing sensitivity. Precise mode is equivalent to *-m 98 -M 98* for TD2.LongOrfs and *-P 0*.*9 --retain-long-orfs-fdr 0*.*005* for TD2.Predict. The *--precise* option must be passed to both TD2.LongOrfs and TD2.Predict.

### False discovery rate control via an extreme value theory of longest ORF length

ORF length is a key feature used to distinguish protein-coding regions from non-coding regions in transcripts. Very long ORFs are unlikely to occur by chance in non-coding RNA, as they require a long stretch of codons unbroken by any stop codon. In most organisms, three out of 64 codons are stop codons: UAA, UGA, and UAG. Thus, in the simplest case, a random codon can be thought of as flipping a biased coin, where *P(tails, i*.*e. stop codon)* = 3/64 and *P(heads, i*.*e. not stop codon)* = 61/64. This is, of course, affected by GC-content and Markov effects, but for the sake of simple explanation we will assume these probabilities to be correct and nucleotides to be independent and identically distributed (i.i.d.).

Rather than calculate the probability of an ORF, defined as a sequence bounded by both a start and a stop codon, we instead calculate the length of the longest stopless sequence (LSS). Disregarding start codons greatly simplifies calculations and leads to similarly useful results. By definition, the LSS length must be greater than or equal to the longest ORF length. The LSS length can be calculated by a geometric distribution, similar to the number of coin tosses in a row without a tails flip.

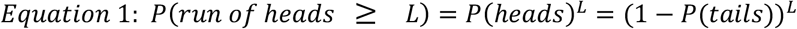

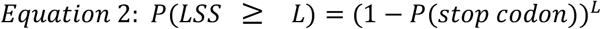

The geometric solution shown in Equation 2 is commonly used to estimate the probability of observing an LSS of a certain length, thus setting a threshold for L that controls the false discovery rate (FDR). However, the geometric solution may be suboptimal for estimating LSS length.

A transcript can be thought of as a sequence of coin flips, where the longest ORF is the longest unbroken run of heads anywhere in the sequence. As such, the solution for the expected value of the longest run of heads, as well as the variance and distribution type, can be derived from extreme value theory. Fortunately, we can stand on the shoulders of giants here: see Gordon, Shilling, and Waterman 1986 ^5^, as well as Schilling 1990 ^6^. Equations 3 and 4 below show the asymptotic expected value and variance of the longest run of heads in a sequence (i.e. the LSS in a transcript) where *n* is the length of the whole sequence (i.e. the length of the transcript), *p* is the probability of heads (i.e. not a stop codon), *q* is the probability of tails (i.e. a stop codon), *γ* is Euler’s constant, *π* is pi, and *r*_*1*_, *ε*_*1*_, *r*_*2*_, and *ε*_*2*_ are periodic functions that quickly tend to zero and, for practical purposes, can be approximated by zero. This “longest run of heads” solution to LSS, with adjustments of *p* and *q* for GC content, can be used to more accurately predict the length of the LSS in a noncoding transcript.

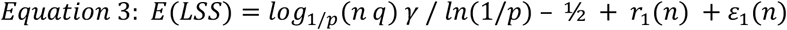

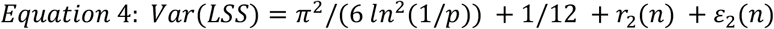

The distribution of LSS lengths across transcripts can be fit by a type-I generalized extreme value distribution, a Gumbel distribution, where μ is the mode and β is a scale parameter. The cumulative distribution function of LSS lengths is defined in Equation 5. Equation 6 and Equation 7 show properties of the Gumbel distribution used to derive the mode μ and shape parameter β.

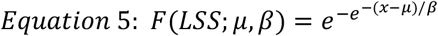

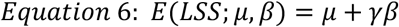

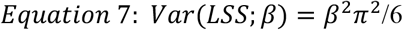

Given Equations 3 through 7, we can calculate the inverse of the cumulative distribution function to find a length *L* that controls the FDR for the LSS in transcripts, adjusted to consider the single longest expected LSS among all six reading frames. The full equation used by TD2 is shown in Equation 8. *L* is the length threshold, in nucleotides, above which we would expect to observe LSS length greater than *L* in fewer than *fdr* as a proportion of the total number of transcripts. *p* is the probability of not getting a stop codon in a random selection of three nucleotides, which varies with GC content. *fdr* is the false discovery rate adjusted for all six frames assuming i.i.d. codon frequency between frames.

In summary, L is the length above which an ORF should be predicted to encode a protein, given transcript length, stop codon likelihood, and a desired false discovery rate in random sequence.

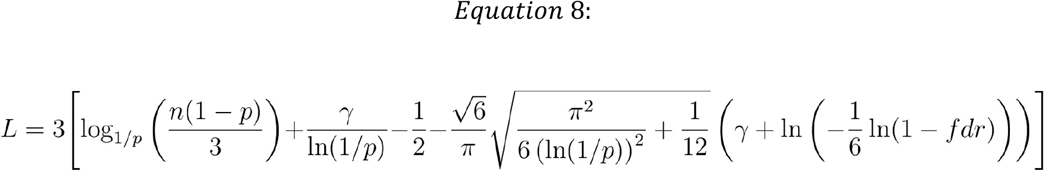

TD2 calculates *L* by default for each transcript, and *fdr* can be modified using the *retain-long-orfs-fdr* parameter. Our implementation can be found as the “*length_at_fdr*” function in TD2.Predict. An analysis of this FDR control technique shows strong concordance with distributions of LSS length in simulated nucleotide sequences. However, this underestimates the FDR of ORFs in human non-coding sequence, in part due to unaccounted-for Markov effects (data not shown). This result may be explored in future work. Thus, FDR parameters in TD2 can be used to limit false discoveries but FDR parameters should not be used as a precise measure of the true FDR in biological data.

The extreme value solution appears to be an improvement for controlling false discovery of long ORFs in transcriptome assemblies, as demonstrated by dramatically reduced false positives in precise mode shown in Table 2 and Table 3 below.

**Table 1:** Performance on high-quality reference transcripts.

|  | # Proteins | TD2 (default) |  | TransDecoder |  | GeneMarkS-T |  |
| --- | --- | --- | --- | --- | --- | --- | --- |
|  |  | TP | FP | TP | FP | TP | FP |
| MANE (human) | 19,354 | **18,771**<br>(97.0%) | **211**<br>(1.1%) | 18,642<br>(96.3%) | 305<br>(1.6%) | 18,762<br>(96.9%) | 287<br>(1.5%) |
| *C. elegans* | 28,589 | **26,627**<br>(93.1%) | 348<br>(1.2%) | 26,185<br>(91.6%) | **230**<br>(0.8%) | 26,335<br>(92.1%) | 257<br>(0.9%) |
| *A. thaliana* | 48,265 | **45,762**<br>(94.8%) | 406<br>(0.8%) | 45,075<br>(93.4%) | 400<br>(0.8%) | 45,300<br>(93.9%) | **383**<br>(0.8%) |
| *P. falciparum* | 5,354 | **5,229**<br>(97.7%) | **2**<br>(0.04%) | 5,174<br>(96.6%) | 5<br>(0.09%) | 5,182<br>(96.8%) | **2**<br>(0.04%) |
| *P. vivax* | 5,392 | **5,216**<br>(96.7%) | 50<br>(0.9%) | 5,147<br>(95.5%) | 42<br>(0.8%) | 5,195<br>(96.3%) | **38**<br>(0.7%) |

**Table 2:** Protein predictions on tuatara transcriptome shotgun assembly.

|  | TD2 (default) | TD2 (*--precise*) | TransDecoder | GeneMarkS-T |
| --- | --- | --- | --- | --- |
| True Positives | **24,068** | 22,181 | 23,084 | 23,201 |
| Complete Proteins | **1,925** | 1,852 | 1,444 | 1,455 |
| Non-matching Predictions | 13,009 | **5,615** | 12,709 | 12,592 |

**Table 3:** Protein predictions on a preliminary assembly of 10,000 GTEx RNA-seq datasets built while creating the CHESS human transcriptome.

|  | TD2 (default) | TD2 (--precise) | TransDecoder | GeneMarkS-T |
| --- | --- | --- | --- | --- |
| True Positives | **322,831** | 302,074 | 304,110 | 306,456 |
| Complete Proteins | **133,775** | 128,843 | 119,312 | 98,880 |
| Non-matching Predictions | 161,051 | **51,656** | 221,844 | 275,534 |

### Tool comparison on model organism transcriptomes

TD2, TransDecoder ^7^, and GeneMarkS-T ^8^ were run with default parameters on high-quality reference transcriptomes including MANE v1.4 (Matched Annotation from NCBI and EMBL-EBI) ^9^, *C. elegans* (GCF_000002985.6) ^10^, *A. thaliana* (GCF_000001735.4) ^11^, *P. falciparum* (GCF_000002765.6) ^12^, and *P. vivax* (GCF_000002415.2) ^13^. No homology search was performed. Tools were run to predict the single best ORF per transcript. True positives were defined generously, requiring an exact match only to the final twenty 3’ amino acids in the reference protein, excluding the stop codon.

TD2’s default settings are intended to maximize sensitivity to proteins while keeping a reasonable cap on false positives. Overall, TD2 with default settings had higher sensitivity than TransDecoder in MANE (human), worm (*C. elegans*), thale cress (*A. thaliana*), and the malaria-causing parasites *P. falciparum* and *P. vivax*. Notably, *P. falciparum* is an extreme outlier in GC-content among eukaryotes.

The RefSeq assembly of *P. falciparum* used here has a GC-content of only 19.34%, relative to 42.9% for *P. vivax*. Because non-protein-coding long ORFs are less likely to occur by chance in low-GC genomes, simply choosing the longest ORF for each *P. falciparum* transcript results in a sensitivity of 99.8% (5,345 proteins). Thus, a “longest ORF” strategy would outperform all tools on *P. falciparum* but would not work well in higher GC-content organisms. This strategy would also result in many false-positives in non-protein-coding transcripts, which raises the question of what should count as a false positive.

Measuring accuracy only on reference transcripts is fundamentally limited because it assumes all transcripts are protein coding. False positives in such a regime can only be generated by choosing the incorrect ORF on a protein-coding transcript. However, many transcripts do not encode proteins. Thus, a comparison of tools should measure false positives in non-coding transcripts as well. A more realistic estimate for the false positive rate (FPR) is explored in the following sections.

### Analysis of the tuatara transcriptome shotgun assembly

Accuracy on highly curated reference transcriptomes, where each given transcript is guaranteed to contain a known protein, is not the best representation of practical performance. TransDecoder and similar tools are often intended to annotate a *de novo* assembled transcriptome. To compare performance in this scenario, we analyzed a transcriptome shotgun assembly (TSA) produced as part of the effort to sequence the tuatara genome (GGNQ00000000.1) ^14^. This TSA was produced by sequencing blood and early-stage embryos with an Illumina HiSeq 2500. Reads were assembled with Trinity v2.2.0. CD-HIT-EST v4.6.6 ^15^ was used to collapse contigs at 98% identity.

The tuatara TSA was annotated using TD2 with default parameters and with the “--*precise*” option enabled, as well as TransDecoder and GeneMarkS-T using default parameters. Only the single best ORF per transcript was kept for all tools. Proteins were considered true positives if any amino acid 30-mer matched any protein in the tuatara reference genome annotation (ASM311381v1). Proteins were considered “Complete Proteins” if they matched at 100% length and 100% identity to annotated proteins in the tuatara reference genome. Parameters for all tools were set to include 3’ partial, 5’ partial, and internal ORFs. It is not possible to prove that an extra prediction is not, in fact, a protein. Some non-matching predictions, i.e. those that do not match any protein in the reference annotation, may be true proteins that were missed by the annotation. However, most non-matching predictions are likely false positives. Thus, tools that are more sensitive while producing fewer overall predictions are preferred.

Depending on the selected mode, TD2 offers either the highest sensitivity or the highest precision on the tuatara TSA. This shotgun assembly represents one use case for TD2, a relatively noisy data set consisting mostly of partial transcripts. In contrast, some transcriptome studies may generate higher-quality, fully assembled transcripts. For this reason, we also tested tools on transcripts from CHESS, a comprehensive dataset of human genes based on nearly 10,000 RNA seq runs from the GTEx project, described in detail by Varabyou et al. 2023 ^16^.

The high-quality published version of CHESS 3 contains 141,100 human transcripts of which 99,201 are protein-coding. Here, we used a superset of CHESS comprising 987,244 noise-filtered transcripts that were assembled from 900 billion reads contained in nearly 10,000 RNA-seq datasets as part of the process of building CHESS ^16^. This large set of transcripts resembles what one might have after a high-quality transcriptome assembly rather than a completed annotation. True positives are determined by an exact amino acid 30-mer to any protein in the reference protein set. Proteins were considered “Complete Proteins” if they matched at 100% length and 100% identity to annotated proteins in the CHESS v3 reference transcript set. If no match is found, predictions are listed as non-matching. Results of all tools on the CHESS preliminary assembly of human transcripts are shown in Table 3.

From this set of 987,244 assembled transcripts, TD2 finds the highest number of ORFs matching proteins in the final CHESS annotation. TD2 also predicts many fewer putative false positives than other tools in both default and precise modes. Note that if ORFs in different transcripts had a 30-amino acid match to the same annotated protein, all of them were considered matches. TD2 (default) did not find 322,831 unique proteins. Rather, it found ORFs with matches to known proteins in 322,831 transcripts. CHESS v3 only contains 73,767 distinct protein sequences in its 99,201 protein-coding transcripts, and many of the matches counted here are duplicates, where two distinct transcripts contain the same ORF, as one might expect in a transcriptome assembly.

### Analysis of false positives from human long non-coding RNA

We used long non-coding RNA (lncRNA) to estimate a false positive rate for each tool in known non-coding sequences. A high-confidence set of human lncRNA was downloaded from LNCipedia version 5.2 ^17^. A total of 107,039 transcripts from 49,372 genes, with all putatively protein-coding genes excluded, were run through TD2, TransDecoder, and GeneMarkS-T. All tools were run to predict at most a single protein per transcript. The number of false positives across a range of minimum ORF lengths are shown in Table 4.

**Table 4:**
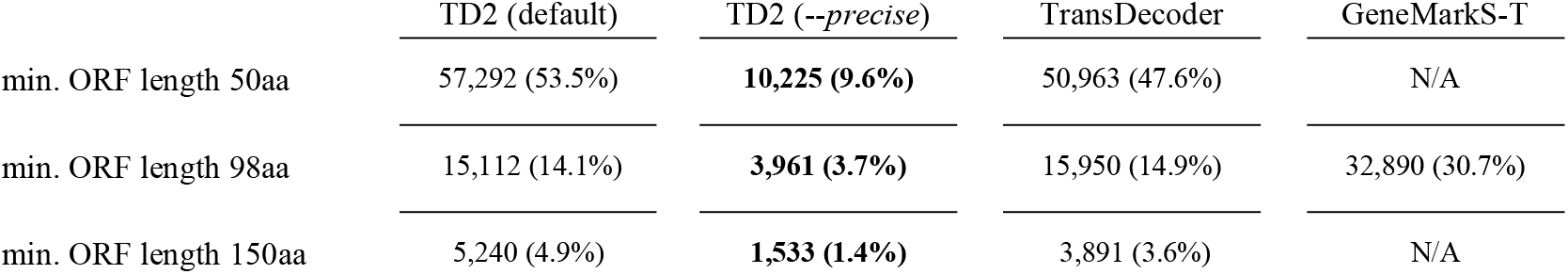
False-positive predictions in human lncRNA.

TD2 achieves the lowest false positive rate (FPR) across a range of minimum ORF length settings. We were not successful in attempts to enable GeneMarkS-T version 5.1 to permit custom minimum ORF lengths. Thus, results are shown only for 98aa, the length it chooses based on the GC content. Of potential interest for future work, TD2 in precise mode maintains reasonable precision even at a minimum ORF length of 50aa, which may enable analysis of short proteins previously unannotated due to high false positive rates at lower ORF lengths.

## CONCLUSIONS

TD2 is a compelling new computational tool for users wishing to annotate transcriptome assemblies with high sensitivity and precision. When non-coding RNA is included, as would be expected in most eukaryotic transcriptomic data, we see a dramatic reduction in the number of putative false positives produced by TD2 relative to other tools.

## AVAILABILITY AND REQUIREMENTS

**Project name**

TD2

**Project home page**

https://github.com/Markusjsommer/TD2

**Operating system**

Linux, Unix/macOS

**Programming language**

Python

**Other requirements**

Pip, PyTorch

**License**

MIT

### Any restrictions to use by non-academics

Permission is hereby granted, free of charge, to any person obtaining a copy of this software and associated documentation files (the “Software”), to deal in the Software without restriction, including without limitation the rights to use, copy, modify, merge, publish, distribute, sublicense, and/or sell copies of the Software, and to permit persons to whom the Software is furnished to do so, subject to the following conditions:

The above copyright notice and this permission notice shall be included in all copies or substantial portions of the Software.

The software is provided “as is”, without warranty of any kind, express or implied, including but not limited to the warranties of merchantability, fitness for a particular purpose and noninfringement. In no event shall the authors or copyright holders be liable for any claim, damages or other liability, whether in an action of contract, tort or otherwise, arising from, out of or in connection with the software or the use or other dealings in the software

## DECLARATIONS

### Availability of data and materials

https://github.com/Markusjsommer/TD2

### Competing interests

The authors declare no conflicts of interest.

## Funding

This work was supported in part by the National Institutes of Health [R01-HG006677, R35-GM0130151]

## Author contributions, CRediT

**MJS:** Conceptualization, Methodology, Software, Validation, Formal analysis, Investigation, Resources, Data Curation, Writing - Original Draft, Writing - Review & Editing, Visualization, Supervision, Project administration. **AM:** Methodology, Software, Validation, Writing - Original Draft, Writing - Review & Editing. **HJJ:** Validation, Writing - Review & Editing. **BJH:** Software, Validation, Writing - Review & Editing. **SLS:** Validation, Writing - Review & Editing, Funding acquisition

## Acknowledgements

We would like to thank all two- and four-legged members of the Salzberg and Pertea labs as well as those residing on Bioinformatics Alley at NBACC and Dr. Michael D. Lee for helpful discussions. We would also like to thank Louis Gordon, Mark F. Schilling, and Michael S. Waterman for their work analysis of long head runs, and Mark F. Schilling in particular for his subsequent work making such analysis more approachable.

## Notes

https://github.com/Markusjsommer/TD2

### Competing Interest Statement

The authors have declared no competing interest.

## REFERENCES

1. Sommer, M. J., Zimin, A. V. & Salzberg, S. L. PSAURON: a tool for assessing protein annotation across a broad range of species. NAR Genom. Bioinform. 7, lqae189 (2025).

2. Altschul, S. F., Gish, W., Miller, W., Myers, E. W. & Lipman, D. J. Basic local alignment search tool. J. Mol. Biol. 215, 403–410 (1990).

3. Finn, R. D., Clements, J. & Eddy, S. R. HMMER web server: interactive sequence similarity searching. Nucleic Acids Res. 39, W29–37 (2011).

4. Steinegger, M. & Söding, J. MMseqs2 enables sensitive protein sequence searching for the analysis of massive data sets. Nat. Biotechnol. 35, 1026–1028 (2017).

5. Gordon, L., Schilling, M. F. & Waterman, M. S. An extreme value theory for long head runs. Probab. Theory Relat. Fields 72, 279–287 (1986).

6. Schilling, M. The longest run of heads. College Mathematics Journal 21, 196–207 (1990).

7. Haas, B. J. et al. De novo transcript sequence reconstruction from RNA-seq using the Trinity platform for reference generation and analysis. Nat. Protoc. 8, 1494–1512 (2013).

8. Tang, S., Lomsadze, A. & Borodovsky, M. Identification of protein coding regions in RNA transcripts. Nucleic Acids Res. 43, e78 (2015).

9. Morales, J. et al. A joint NCBI and EMBL-EBI transcript set for clinical genomics and research. Nature 1–6 (2022).

10. The C. elegans Sequencing Consortium*. Genome sequence of the nematode C. elegans : A platform for investigating biology. Science 282, 2012–2018 (1998).

11. Lamesch, P. et al. The Arabidopsis Information Resource (TAIR): improved gene annotation and new tools. Nucleic Acids Res. 40, D1202–10 (2012).

12. Gardner, M. J. et al. Genome sequence of the human malaria parasite Plasmodium falciparum. Nature 419, 498–511 (2002).

13. Carlton, J. M. et al. Comparative genomics of the neglected human malaria parasite Plasmodium vivax. Nature 455, 757–763 (2008).

14. Gemmell, N. J. et al. The tuatara genome reveals ancient features of amniote evolution. Nature 584, 403–409 (2020).

15. Fu, L., Niu, B., Zhu, Z., Wu, S. & Li, W. CD-HIT: accelerated for clustering the next-generation sequencing data. Bioinformatics 28, 3150–3152 (2012).

16. Varabyou, A. et al. CHESS 3: an improved, comprehensive catalog of human genes and transcripts based on large-scale expression data, phylogenetic analysis, and protein structure. Genome Biol. 24, 249 (2023).

17. Volders, P.-J. et al. LNCipedia: a database for annotated human lncRNA transcript sequences and structures. Nucleic Acids Res. 41, D246–51 (2013).

